## Supplementary information for "Dynamic inter-domain transformations mediate the allosteric regulation of human 5, 10-methylenetetrahydrofolate reductase"

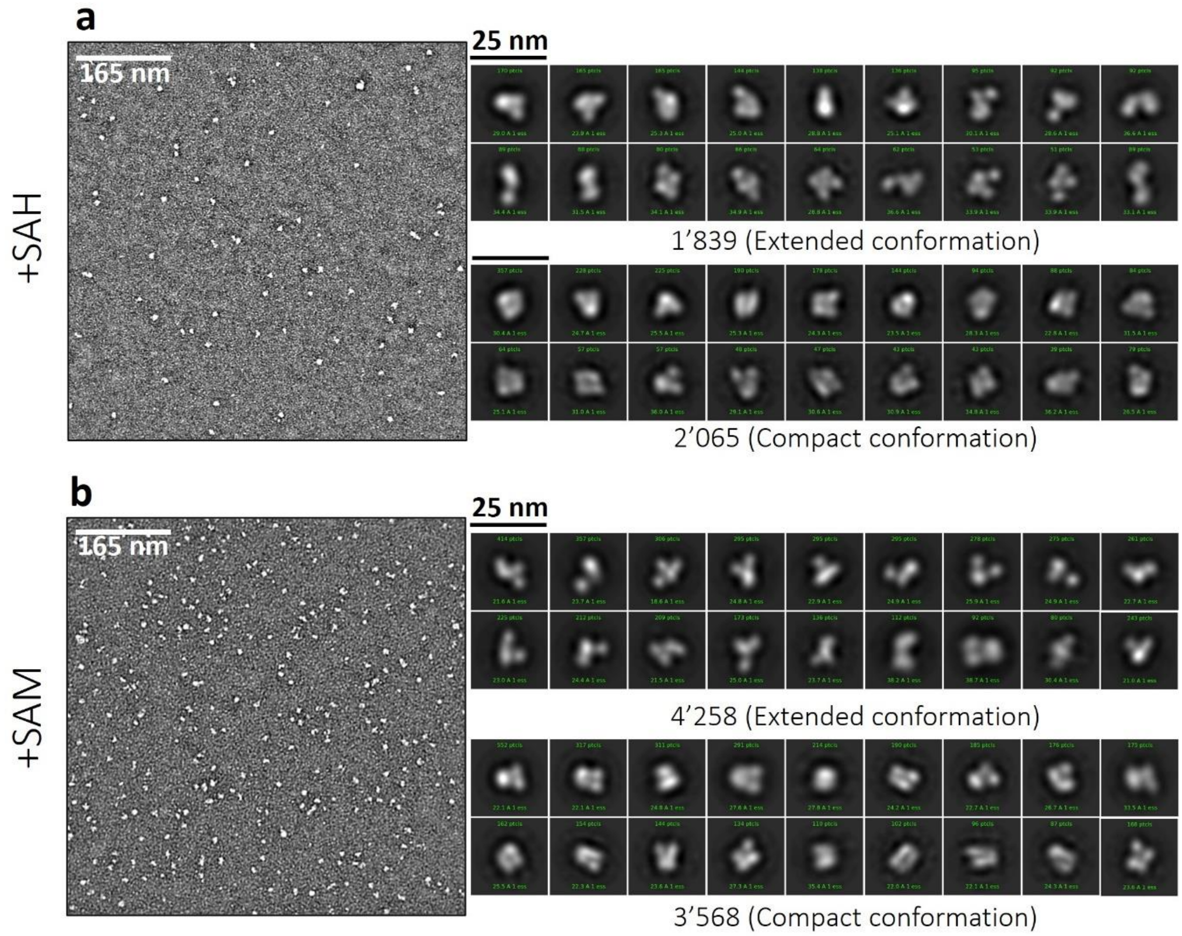


**Supplementary Fig. 1 | Negative stain**. Representative micrographs and 2D classes for **a**, $MTHFR_{FL}$ incubated with 2.6 mM SAH and **b**, $MTHFR_{FL}$ incubated with 5 mM SAM. Both samples exhibit two conformations: one more extended conformation (upper rows) resembling the structure of $MTHFR_{trunc}^{SAH}$ (PDB:6fcx[1]) and one more compact conformation (lower rows) likely to represent inhibited MTHFR.

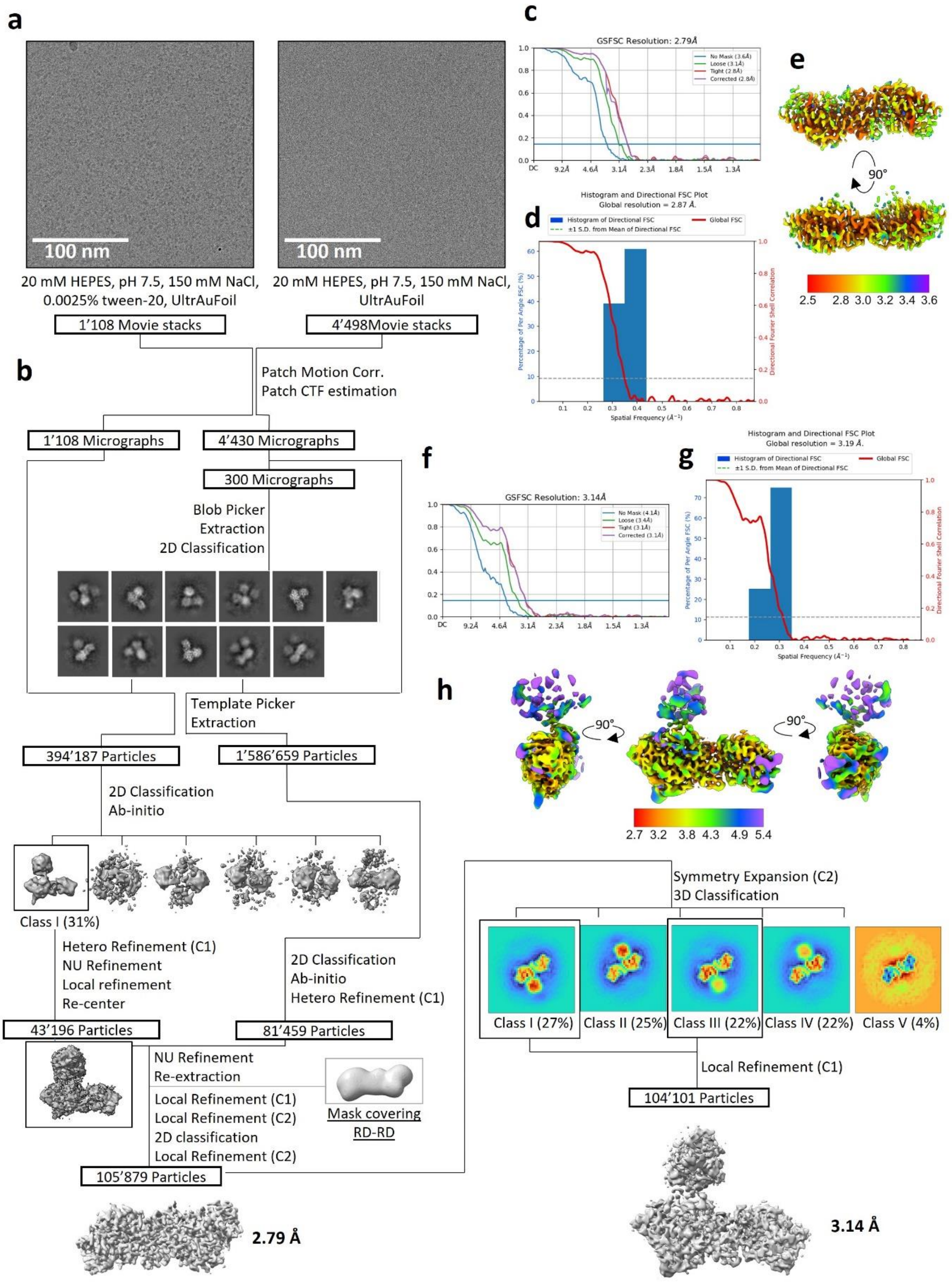


**Supplementary Fig. 2 | Cryo-EM image processing workflow of MTHFR$_{FL}^{SAH}$.** **a**, Representative micrographs from two movie stacks collected with different grid conditions. **b**, Process flow chart for pooled data sets from which two maps for MTHFR$_{FL}^{SAH}$ were reconstructed: One symmetric map MTHFR$_{FL}^{SAH\ (symm)}$ capturing the two regulatory domains at 2.79 Å with corresponding **c**, FSC curve **d**, 3D-FSC plot showing the angular distribution and **e**, local resolution distribution. One asymmetric map

$MTHFR_{FL}^{SAH\ (asymm)}$ capturing the two regulatory domains and low-resolution volume, due to high flexibility, representing one catalytic domain and linker region at average resolution of 3.14 Å, with corresponding **f**, FSC curve **g**, 3D-FSC plot showing the angular distribution and **h**, local resolution distribution.

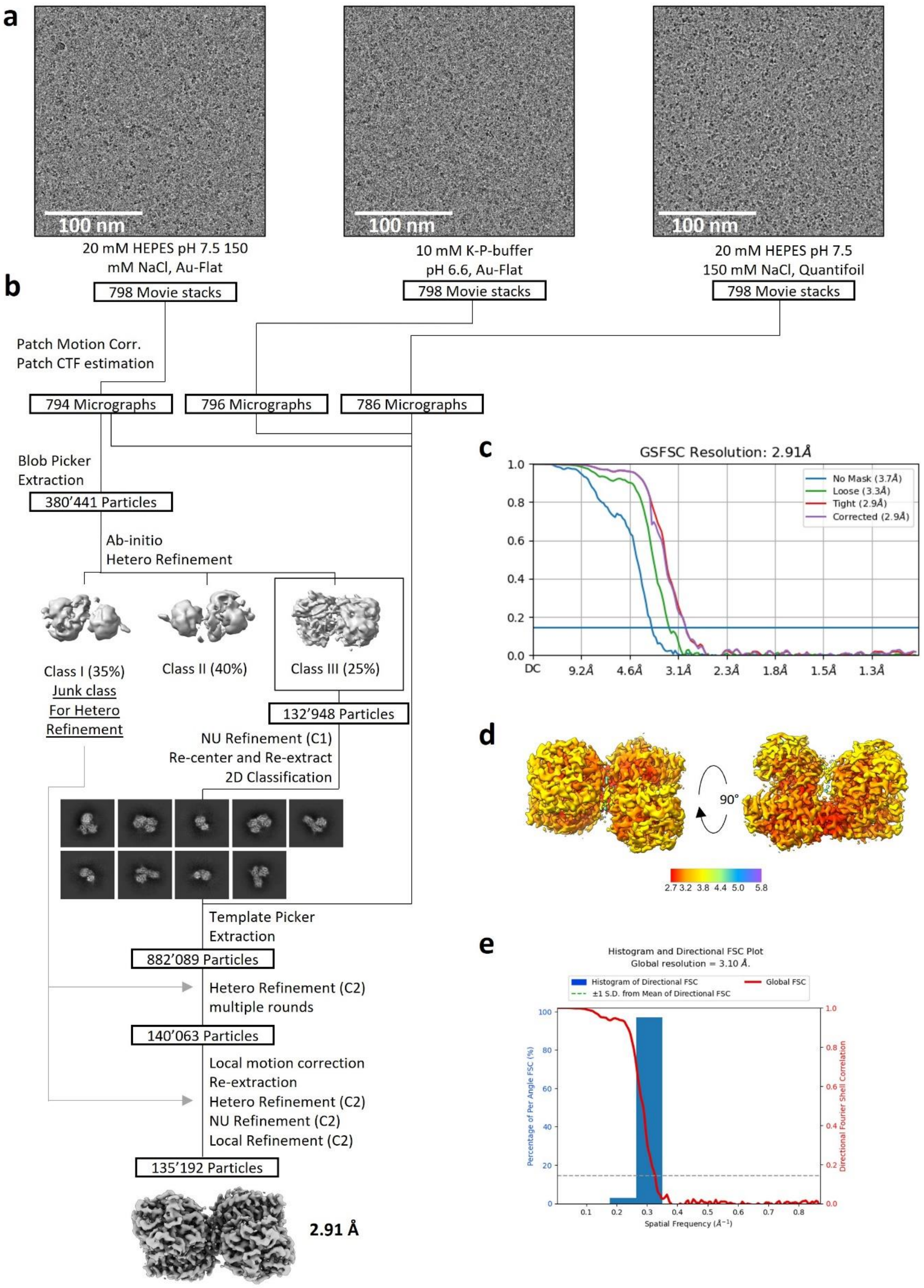


**Supplementary Fig. 3 | Cryo-EM image processing workflow of MTHFR$_{FL}^{SAM}$. a**, Representative micrographs from three movie stacks collected with different grid conditions. **b**, Process flow chart for pooled data sets from which one map for $MTHFR_{FL}^{SAM}$ at 2.91 Å was reconstructed with corresponding **c**, FSC curve **d**, local resolution distribution. and **e**, 3D-FSC plot showing the angular distribution.

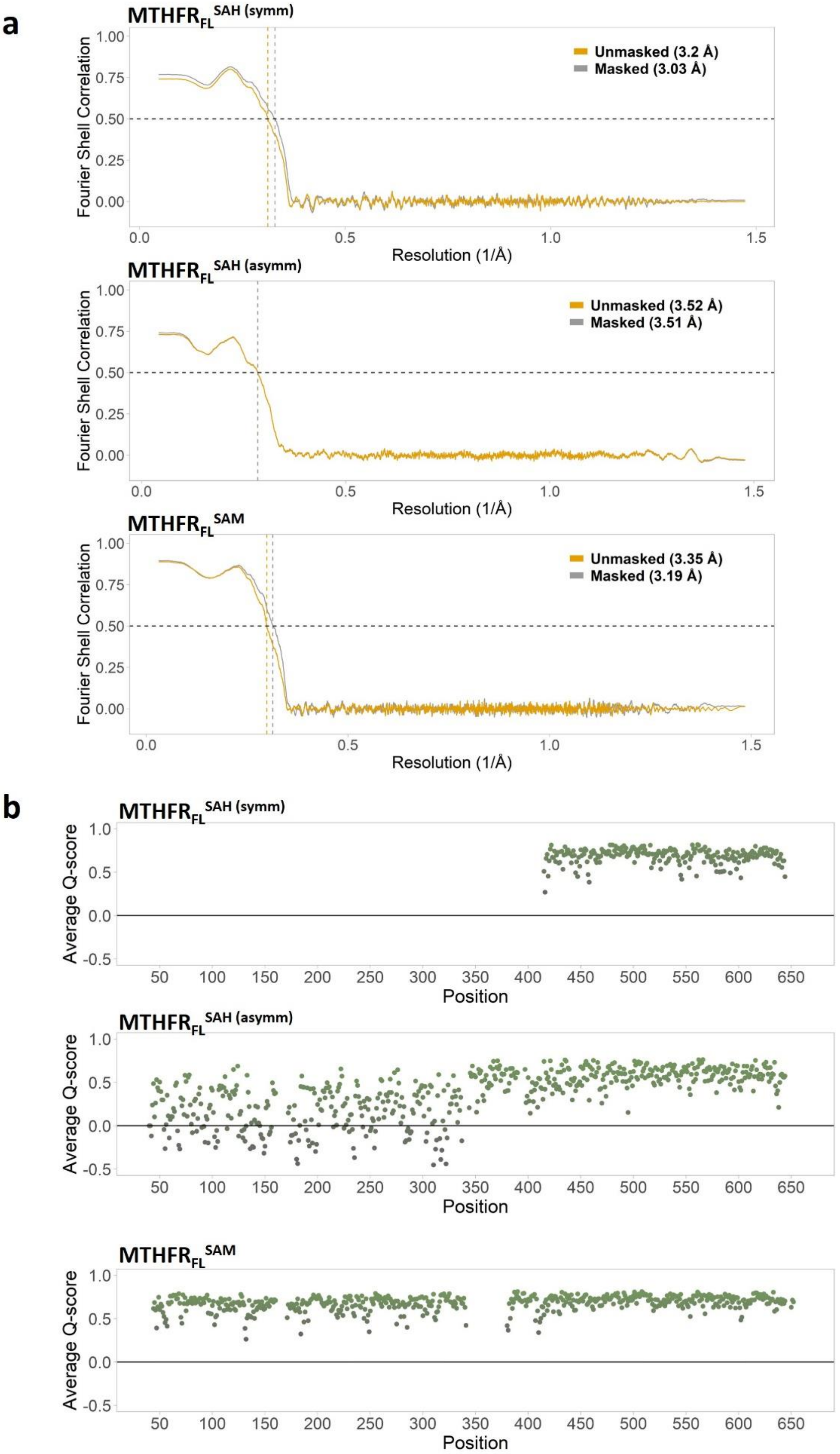


**Supplementary Fig. 4 | Map and model quality of the MTHFR structures. a**, Map to model FSC curves of the A-chains in MTHFR$_{FL}^{SAH\ (symm)}$, MTHFR$_{FL}^{SAH\ (asymm)}$, and MTHFR$_{FL}^{SAM}$ as determined by Phenix validation[2] with printed resolution at FSC=0.5. **b**, Average Q-score[3] for each amino acid position in the

A-chains in $MTHFR_{FL}^{SAH\ (symm)}$, $MTHFR_{FL}^{SAH\ (asymm)}$, and $MTHFR_{FL}^{SAM}$. The low scores for residues 48 to 338 of $MTHFR_{FL}^{SAH\ (asymm)}$ are due to the low resolution of this region corresponding to the flexible CD.

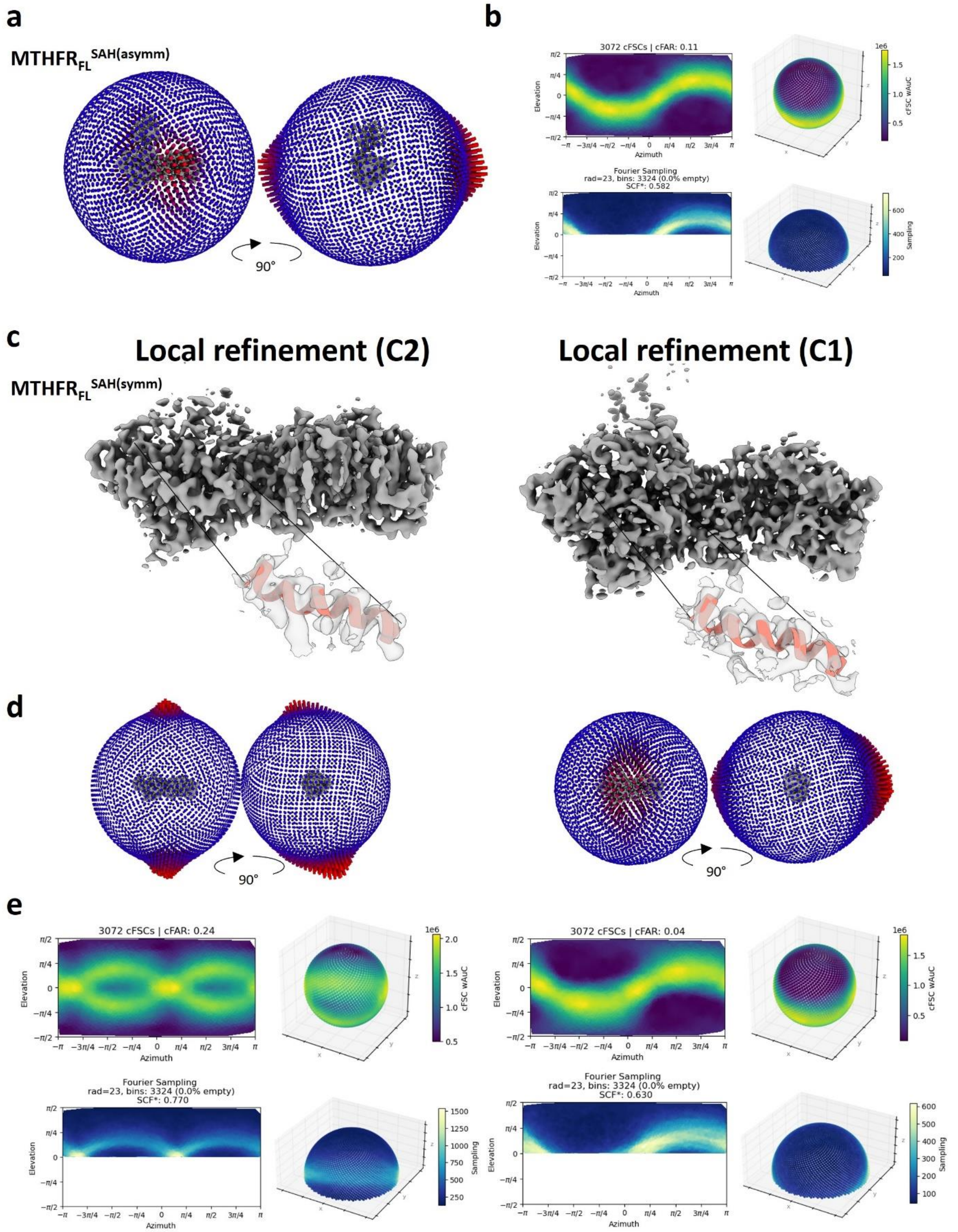


**Supplementary Fig. 5 | Evaluation of C1 and C2 symmetry of MTHFR$_{FL}^{SAH}$ processing. a**, 3D viewing direction distributions and **b**, cFAR and SCF* values for final map for MTHFR$_{FL}^{SAH(asymm)}$. Masked local refinement of the central regulatory domain dimer of the MTHFR SAH bound sample with C2 (deposited map MTHFR$_{FL}^{SAH(symm)}$) and without C2 symmetry and using the same mask. **c**, Resulting sharpened maps of masked local refinements of the RD dimer with C2 symmetry, left, and no symmetry applied (C1), right. An alpha-helix representing residues 576-595 is shown fitted in the density for both maps. It is clear from visual inspection that the C2 symmetric map has secondary structure features whereas the locally refined map without symmetry does not. **d**, The corresponding 3D viewing direction distributions and **e**, cFAR and SFC* plots for the C2 symmetric, left, and C1, right, locally refined maps. An improvement in both cFAR and SFC* values are apparent when C2 symmetry is applied however the low cFAR of 0.24 indicates effects from orientation bias remain in the C2 symmetric map.

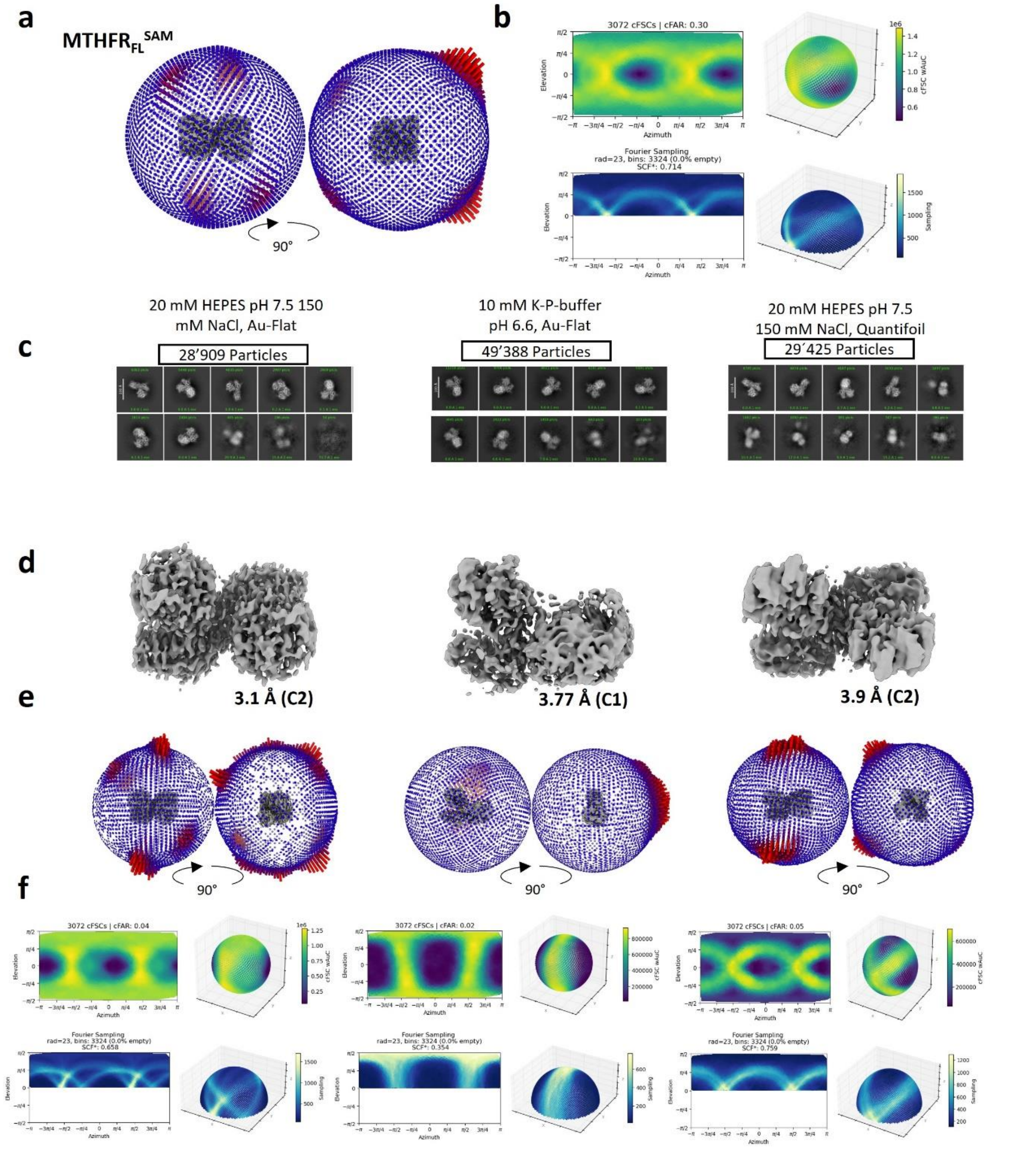


**Supplementary Fig. 6 | Evaluation of the effect of grid conditions on orientation bias**. **a**, 3D viewing direction distributions and **b**, cFAR and SCF* values for final map for $MTHFR_{FL}^{SAM}$. Respective grid conditions pooled to generate $MTHFR_{FL}^{SAM}$ processed individually from template picker job in Supplementary figure 3 with corresponding **c**, 2D classification job, followed by Ab-initio (not shown) and **d**, NU refinement with applied C1 symmetry, and when applicable C2 symmetry. Due to orientation bias grid condition “10 mM K-P-buffer pH 6.6, Au-Flat“ was not able to resolve the C2 symmetry. **e**, 3D viewing direction distributions and **f**, cFAR and SCF* values.

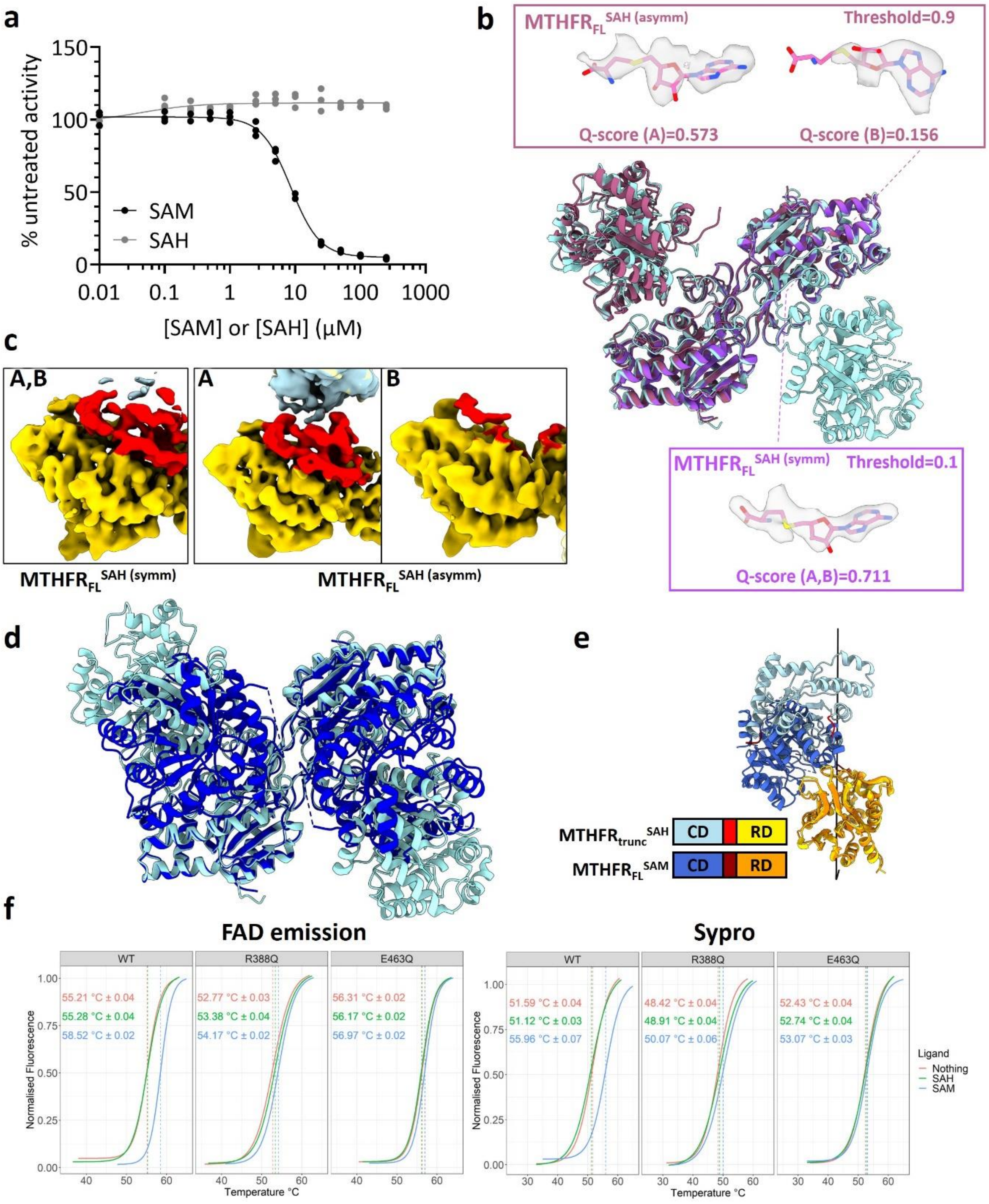


**Supplementary Fig. 7 | Structural and biochemical response upon MTHFR$_{FL}$ binding SAM/SAH. a**, Relative enzymatic activity of purified recombinant MTHFR$_{FL}$ following incubation with SAM or SAH. N=3 technical replicates. **b**, Cryo-EM models MTHFR$_{FL}^{SAH\ (asymm)}$ (dark pink) (Cα-RMSD 0.826 Å) and MTHFR$_{FL}^{SAH\ (symm)}$ (purple) (C$^{\alpha}$-RMSD 0.536 Å) superimposed on MTHFR$_{trunc}^{SAH}$ (light blue), aligned using the homodimeric RD interface. Inset shows electron densities for the sharpened map at specified threshold and range of 2.2 Å. The average Q-score is also shown for each SAH molecule in their respective protomers (A, B). **c,** Zoomed view of the linker (red) with surrounding regulatory domain (yellow) and catalytic domain (blue) in respective protomer (A, B) of the cryo-EM maps MTHFR$_{FL}^{SAH\ (symm)}$ (left) and MTHFR$_{FL}^{SAH\ (asymm)}$ (middle and right) **d,** Cryo-EM model MTHFR$_{FL}^{SAM}$ (dark blue) (C$^{\alpha}$-

RMSD 0.691 Å) superimposed on the $MTHFR_{trunc}^{SAH}$ crystal structure (light blue). **e**, Hinge axis generated with DynDom[4] comparing the relative position of the catalytic domain between $MTHFR_{FL}^{SAM}$ (dark blue) and $MTHFR_{trunc}^{SAH}$ (light blue). **f**, Differential fluorimetry of FAD emission (left) or Sypro dye (right) of purified recombinant $MTHFR_{FL}$ or protein variants following incubation with 1 mM SAM, 1 mM SAH or no ligand. Tm calculated from inflation point was estimated using a Boltzmann curve fit function in R. N=6 technical replicates.

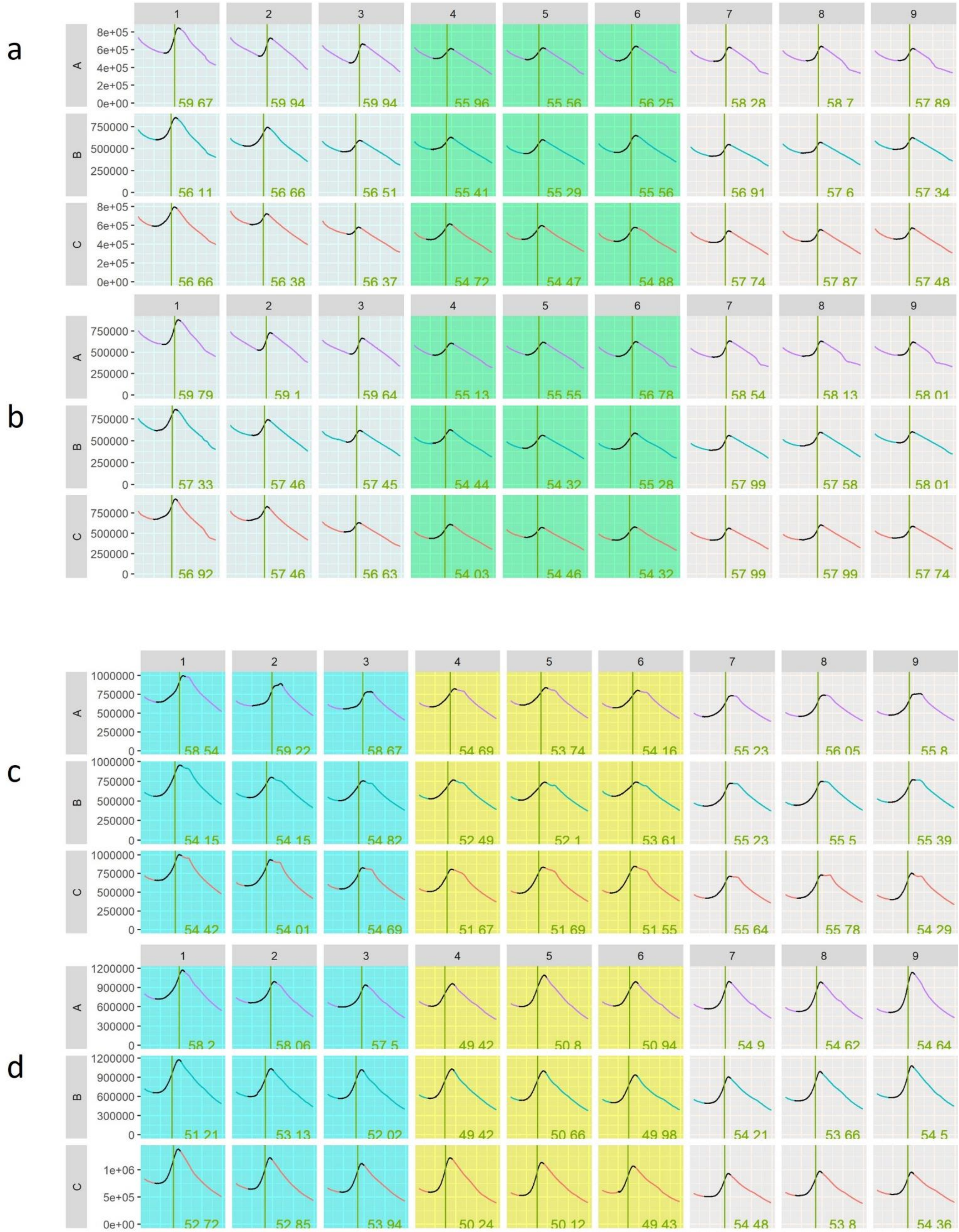


**Supplementary Fig. 8 | Raw data for Differential Scanning Fluorimentry.** Raw data curves of Fluorescence (y-axis) versus temperature (x-axis) as given from QuantStudio™ 7 Pro and visualised with R. Black curve line represents data which were later normalised and used to derive Tm using

Boltzmann curve fit in R. The temperature range for the black curve was defined by data presenting a positive first derivative surrounding the maximum inclination point (green line and number) as calculated from the raw derivative given by QuantStudio™ 7 Pro. The temperature range was then extended by ±2 degrees. SAM (Rows: A), SAH (Rows: B) and Nothing (Rows: C), $MTHFR_{FL}$ (Column 1,2,3), Arg388Gln (Column 4,5,6) and Glu463Gln (Column 7,8,9). Each condition was measured with N=6 technical replicates. **a,b** Measurements utilising FAD quenching. **c,d** Measurements utilising emission from Sypro orange.

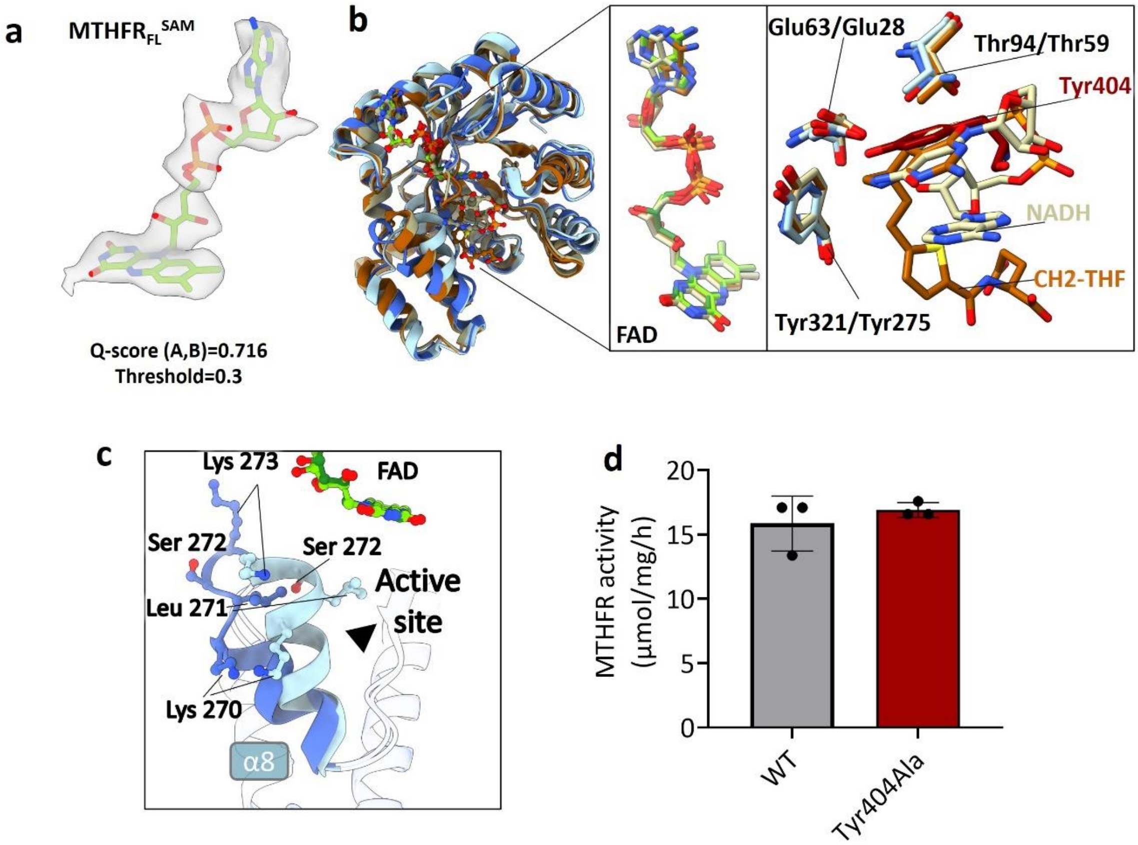


**Supplementary Fig. 9 | The catalytic domain accommodates an auto-inhibitory element while maintaining a conserved structural integrity. a**, Electron density for the sharpened map at specified threshold, and range of 2.2 Å for FAD found in the respective protomer (A, B) of $MTHFR_{FL}^{SAM}$. The average Q-score is also shown. **b**, Superimposition of human $MTHFR_{trunc}^{SAH}$ (light blue), human $MTHFR_{FL}^{SAM}$ (dark blue), *Escherichia coli* MTHFR co-crystallised with NADH (PDB:1zrq, beige), and *Escherichia coli* MTHFR with Ala177Val background co-crystalized with 5,10-dideazafolate analogue (LY309887) representing the position of 5,10-methylenetetrahydrofolate ($CH_2$-THF) (PDB:2fmn, brown). Inset showing a zoomed view of FAD (left) and residues interacting with Tyr404 (dark red) within the active site. **c**, [55,56]Zoomed view of superimposed $MTHFR_{trunc}^{SAH}$ (light blue) and $MTHFR_{FL}^{SAM}$ (dark blue) showing the movement of alpha helix 8 (α8) to accommodate hydrophobic linker insertion in the active site. **d**, Specific activity of overexpressed $MTHFR_{FL}$ (WT) and alanine substitution of Tyr404. N=3 technical replicates.

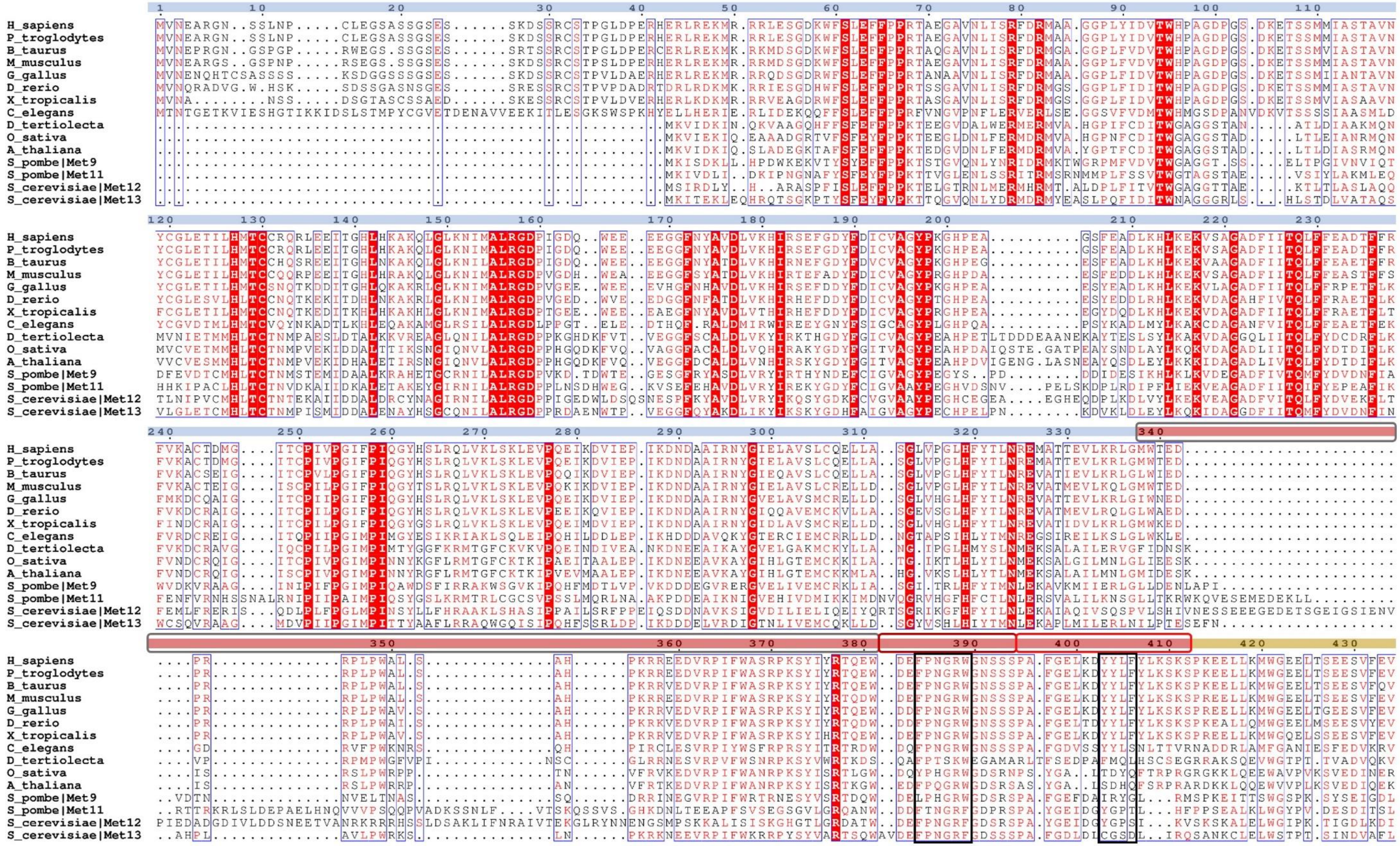
H_sapiens
P_troglodytes
B_taurus
M_musculus
G_gallus
D_rerio
X_tropicalis
C_elegans
D_tertiolecta
O_sativa
A_thaliana
S_pombe|Met9
S_pombe|Met11
S_cerevisiae|Met12
S_cerevisiae|Met13

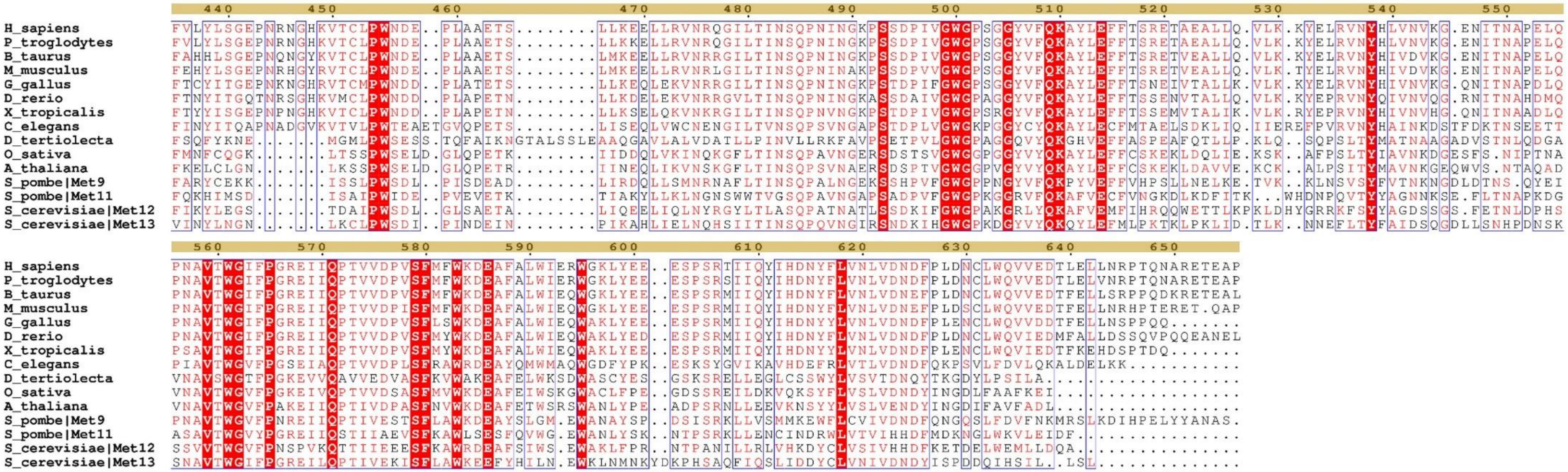


**Supplementary Fig. 10 | Sequence alignment for Eucaryotc MTHFR$_{FL}$.** Sequences were manually selected from MTHFR entries in UniProt (June 2023) and aligned with Clustal Omega, and visualised with ESPript 3.0 (global score: 0.5) [5]. Marked catalytic domain (1-337), linker (338-412) and regulatory domain (413-656). *Organism*-general name (UniProt ID): *Arabidopsis thaliana* - Mouse-ear cress (O80585), *Bos taurus* - Cattle (Q5I598), *Caenorhabditis elegans* - Nematode worm (Q17693*)*, *Danio rerio* - Zebrafish (B0V153), *Dunaliella tertiolecta* - Green alga (A0A7S3VQX8), *Gallus* - Chicken (E1BXL1), *Homo sapiens* - Human (P42898), *Mus musculus* - House mouse (Q9WU20), *Oryza sativa* - Rice (Q75HE6), *Pan troglodytes* - Chimpanzee (H2PY11), *Saccharomyces cerevisiae* - Brewer's yeast (Met12/P46151, Met13/P53128), *Schizosaccharomyces pombe* - Fission yeast (Met9/Q10258, Met11/O74927), *Xenopus tropicalis* - Western clawed frog (A4IIX8). Sequences features marked: Catalytic domain 1-337 (blue shadow), linker 338-412 (red shadow), regulatory domain 413-656(yellow shadow), LS1 (grey box), LS2 (dark red box), LS3 (orange box), $Y_{403}YLF_{406}$ and $F_{384}PNGRW_{389}$ (black boxes).

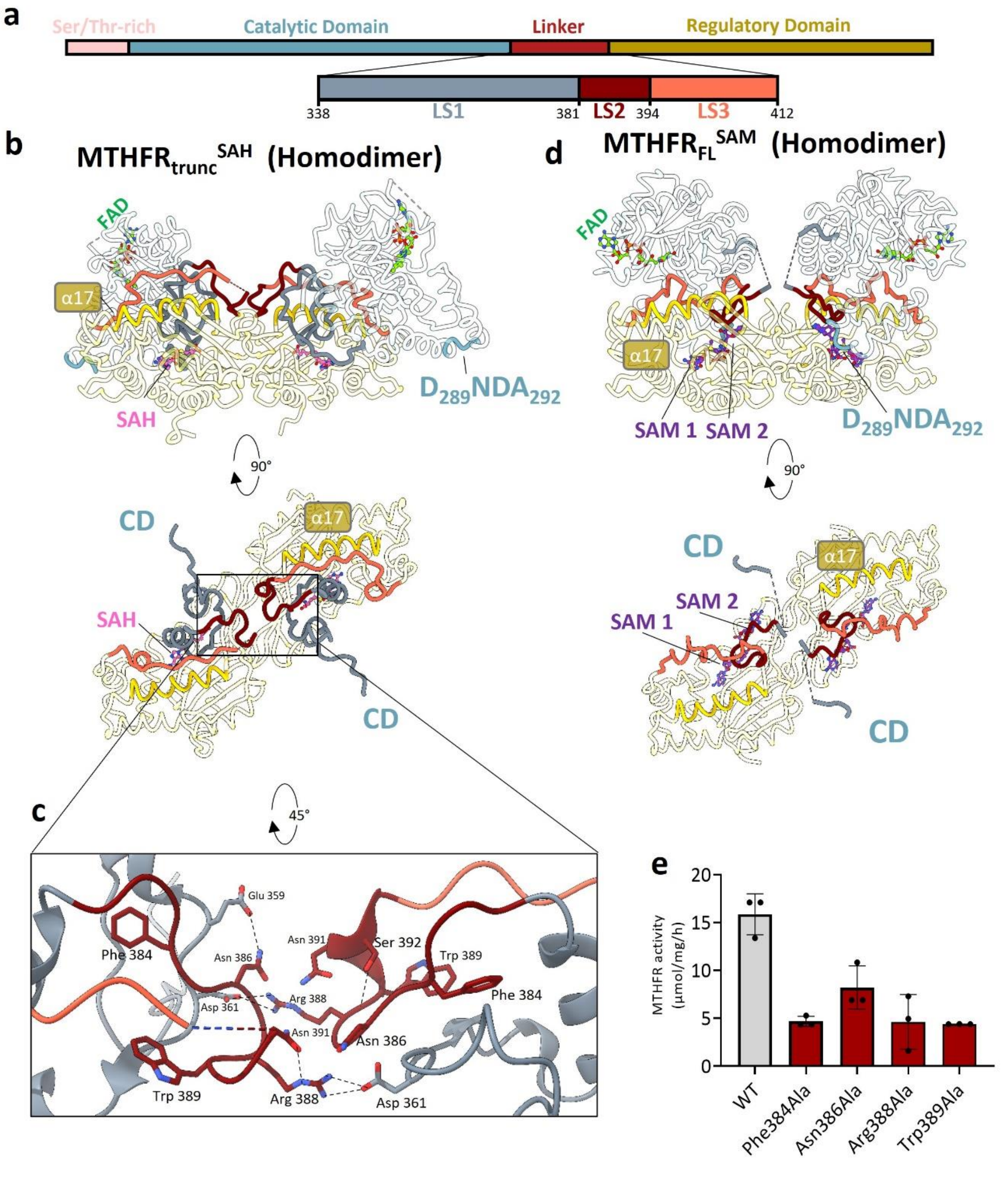


**Supplementary Fig. 11 | Rearrangement of linker segments LS1, LS2 and LS3. a,** Domain diagram of human $MTHFR_{FL}$, coloured according to MTHFR protein domains and with a zoom-in of linker segments LS1 (aa 338-380), LS2 (aa 381-393) and LS3 (aa 394-412). The same colour coding is maintained in subsequent structural representations **b, c, and d.** Transparent models of dis-inhibited $MTHFR_{trunc}^{SAH}$ (**b**) and inhibited $MTHFR_{FL}^{SAM}$ (**d**) with pronounced linker segments LS1, LS2 and LS3, showing their movement in relation to alpha-helix 17 (α17) and CD segment $D_{289}NDA_{292}$. **c**, Inset showing a zoom-in of the homodimeric interface in $MTHFR_{trunc}^{SAH}$, which is further stabilised by interactions between opposing LS2 in the two protomers. **e**, Specific activity data for overexpressed $MTHFR_{FL}$ and alanine substitutions within LS2. N=3 technical replicates.

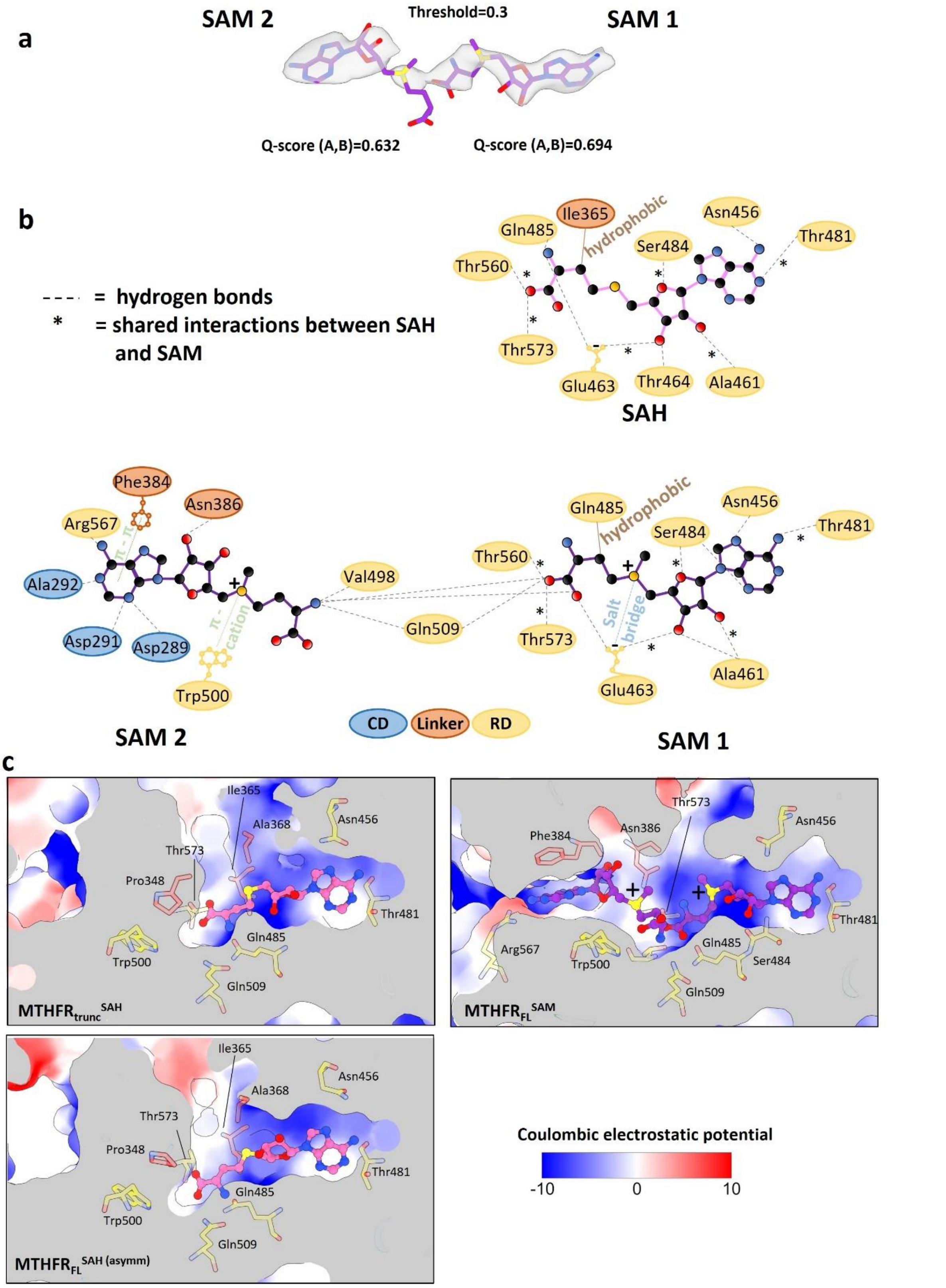


**Supplementary Fig. 12 | Structure and properties of the allosteric pocket. a**, Electron density for sharpened map at specified threshold and range of 2.2 Å, with Q-score for SAM1 and SAM2 in $MTHFR_{FL}^{SAM}$. **b**, Schematic depicting interactions within the allosteric pocket of $MTHFR_{trunc}^{SAH}$ for SAH, and within $MTHFR_{FL}^{SAM}$ for SAM1 and SAM2. Indicated interactions were predicted by PLIP[6], except for the hydrogen bond between SAM1 and SAM2, which were predicted by ChimeraX[7].**c**, Allosteric pocket for $MTHFR_{trunc}^{SAH}$, $MTHFR_{FL}^{SAH}$ and $MTHFR_{FL}^{SAM}$ with coulombic charge predicted with ChimeraX. Negative values in blue and positive in red, showing bound SAH (left) or SAM1 and SAM2 (right).

**a**

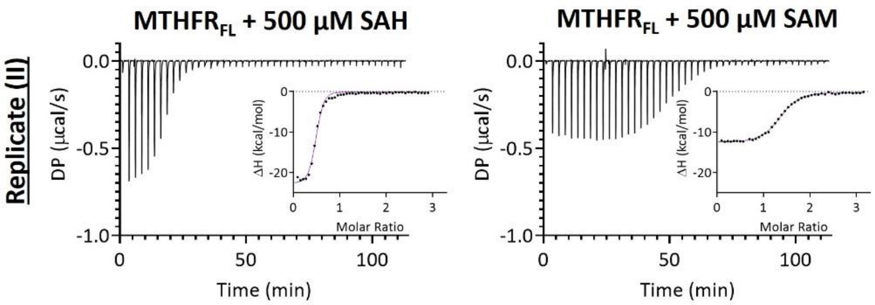


| 45 injections | | | | | | | | |
|---|---|---|---|---|---|---|---|---|
| Sample | N ± error | Average ± S.D. | KD ± error (µM) | Average ± S.D. | ΔH ± error (kcal/mol) | Average ± S.D. | -TΔS (kcal/mol) | Average ± S.D. |
| MTHFR$_{FL}$ + SAM (I) | 1.35 ± 0.00 | 1.33 ± 0.02 | 0.86 ± 0.04 | 0.93 ± 0.07 | -12.80 ± 0.07 | -12.95 ± 0.15 | 4.48 | 4.70 ± 0.22 |
| MTHFR$_{FL}$ + SAM (II) | 1.31 ± 0.01 | | 0.99 ± 0.08 | | -13.10 ± 0.13 | | 4.92 | |
| MTHFR$_{FL}$ + SAH (I) | 0.48 ± 0.00 | 0.47 ± 0.01 | 0.38 ± 0.03 | 0.38 ± 0.00 | -23.70 ± 0.20 | -23.40 ± 0.30 | 15.00 | 14.65 ± 0.35 |
| MTHFR$_{FL}$ + SAH (II) | 0.47 ± 0.00 | | 0.39 ± 0.04 | | -23.10 ± 0.28 | | 14.30 | |

**b**

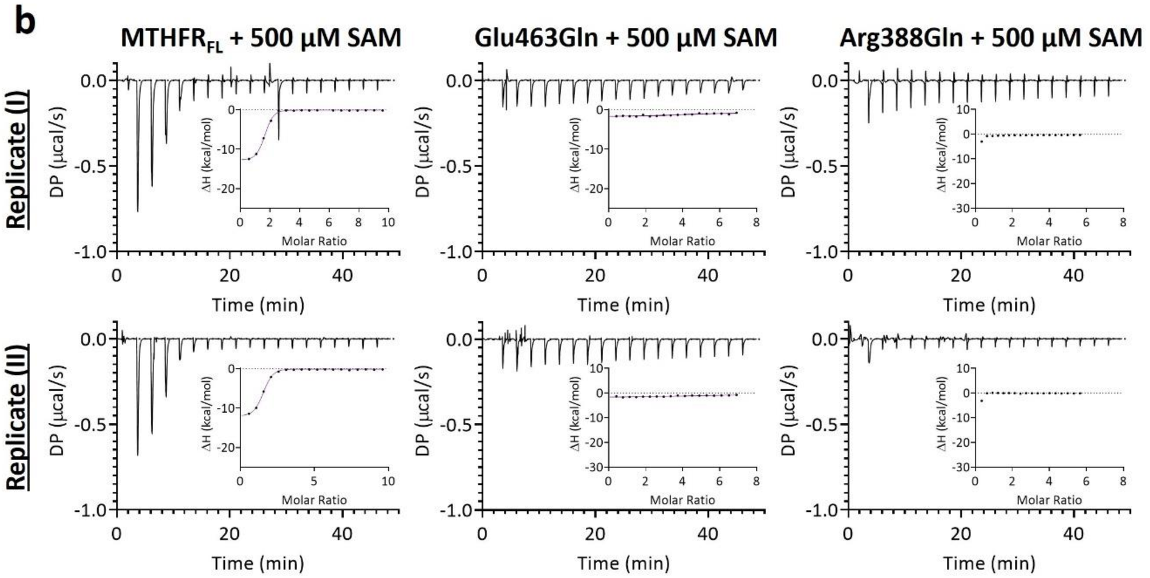


| 19 injections | | | | | | | | |
|---|---|---|---|---|---|---|---|---|
| Sample | N ± error | Average ± S.D. | KD ± error (µM) | Average ± S.D. | ΔH ± error (kcal/mol) | Average ± S.D. | -TΔS (kcal/mol) | Average ± S.D. |
| MTHFR$_{FL}$ + SAM (I) | 1.44 ± 0.02 | 1.40 ± 0.05 | 0.42 ± 0.07 | 0.51 ± 0.09 | -13.00 ± 0.27 | -12.65 ± 0.35 | 4.33 | 4.09 ± 0.24 |
| MTHFR$_{FL}$ + SAM (II) | 1.35 ± 0.02 | | 0.59 ± 0.07 | | -12.30 ± 0.22 | | 3.85 | |
| E463Q + SAM (I) | 7.67 ± 1.05 | 7.74 ± 0.07 | 27.60 ± 32.2 | 26.90 ± 0.70 | -2.13 ± 0.58 | -2.08 ± 0.05 | -4.09 | -4.16 ± 0.07 |
| E463Q + SAM (II) | 7.81 ± 0.97 | | 26.20 ± 26.4 | | -2.02 ± 0.45 | | -4.23 | |
| R388Q + SAM (I) | N/A | N/A | N/A | N/A | N/A | N/A | N/A | N/A |
| R388Q + SAM (II) | N/A | | N/A | | N/A | | N/A | N/A |

**Supplementary Fig. 13 | Isothermal Titration Calorimetry.** Isothermal titration calorimetry (ITC) experiments displaying the heat exchange over time. DP: power differential. The insets illustrate ΔH (change in enthalpy) plotted against the molar ratio. All curves have been normalized against buffer injected into respective MTHFR$_{FL}$ variants. **a,** Above: Shows 45 injections of 500 µM SAM or 500 µM SAH into 30 µM purified recombinant MTHFR$_{FL}$. N=2 technical replicates, where replicate I is shown in Fig. 4a. Below: Table of best-fit parameters (**±** corresponding fitting errors) for each replicate and the mean (**±** S.D.) for each sample condition. **b,** Above: Shows 19 injections of 500 µM SAM in 10 µM purified recombinant $MTHFR_{FL}$ variants. N=2 technical replicates. Below: Table of best-fit parameters (**±** corresponding fitting errors) for each replicate and the mean (**±** S.D.) for each sample condition.

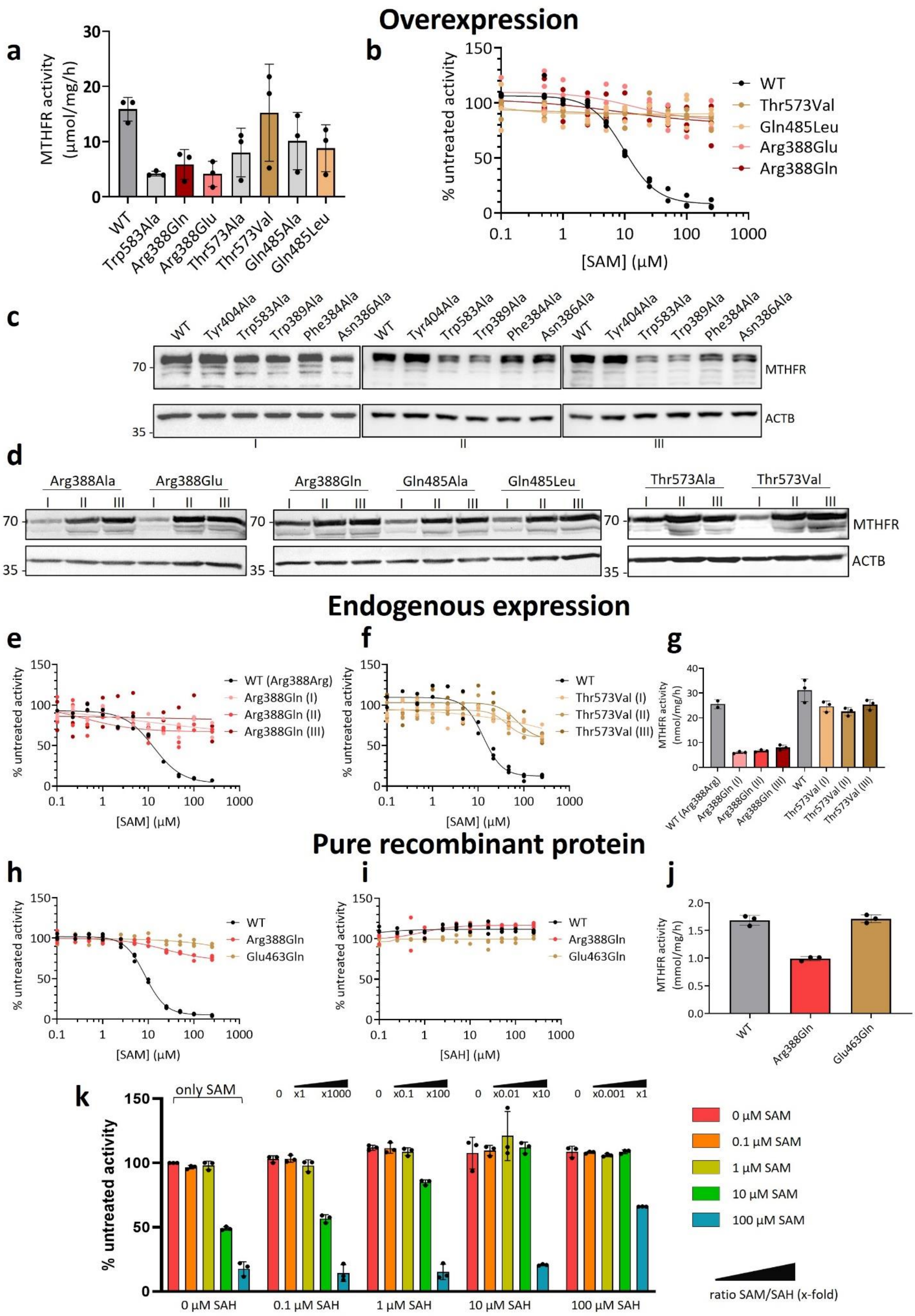
Overexpression
a
b
c
d
Endogenous expression
e
f
g
Pure recombinant protein
h
i
j
k
MTHFR activity (µmol/mg/h)
WT
Trp583Ala
Arg388Gln
Arg388Glu
Thr573Ala
Thr573Val
Gln485Ala
Gln485Leu
% untreated activity
[SAM] (µM)
[SAH] (µM)
Tyr404Ala
Trp389Ala
Phe384Ala
Asn386Ala
MTHFR
ACTB
Arg388Ala
WT (Arg388Arg)
Arg388Gln (I)
Arg388Gln (II)
Arg388Gln (III)
Thr573Val (I)
Thr573Val (II)
Thr573Val (III)
MTHFR activity (nmol/mg/h)
MTHFR activity (mmol/mg/h)
Glu463Gln
only SAM
0 µM SAM
0.1 µM SAM
1 µM SAM
10 µM SAM
100 µM SAM
0 µM SAH
0.1 µM SAH
1 µM SAH
10 µM SAH
100 µM SAH
ratio SAM/SAH (x-fold)

**Supplementary Fig. 14 | Activity data for wild-type (WT) MTHFR and variants.** Specific activity and inhibition curves (WT) MTHFR and protein variants overexpressed in MTHFR knock-out 293T cells (**a-d**), genetically engineered for endogenous expression in 293T cells (**e-g**), or *as purified* recombinant **MTHFR$_{FL}$** protein following SF9 insect expression (**h-k**). **a**, Specific activity. **b**, Inhibition curves for variants Arg388Gln/Glu, Thr573Val and Gln485Leu with WT corresponding to WT seen in Fig. 2e and Fig. 4b,c. Western blotting analysis corresponding to all overexpressed proteins using **c,** 10% acrylamide gel and **d**, 12% acrylamide gel. N=3 biological replicates. **e-f**, Inhibition curves for endogenously expressed WT MTHFR and three clones each for **e**, Arg388Gln, and **f**, Thr573Val, generated with CRISPR/Cas9 gene editing. **g**, Specific activity for each. WT (Arg388Arg) represents a clone which had undergone CRISPR/Cas9 gene editing but showed no mutation within the MTHFR gene; WT represents the parental, unmodified 293T cells. N=3 technical replicates. **h-j**, Inhibition curves for purified recombinant MTHFR$_{FL}$ (WT) and variants Arg388Gln and Glu463Gln with **h,** SAM and **i**, SAH. **j**, Specific activity. N=3 technical replicates. **k,** Competition assay of purified recombinant MTHFR$_{FL}$ (WT) showing SAM inhibition in the presence of increasing concentration of SAH. Samples were pre-incubated with five different concentrations (0, 0.1, 1, 10 and 100 µM) of SAM and SAH respectively. Indication of SAM/SAH ratio are marked out.

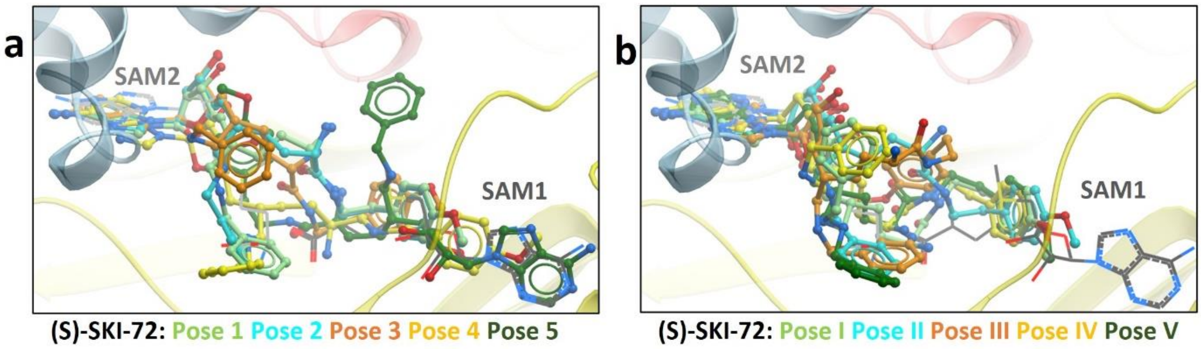


**Supplementary Fig. 15 | Docking poses for (S)-SKI-72 docked into MTHFR structure solved in this work**. Five poses were obtained each from docking performed after superimposing the adenine ring of (S)-SKI-72 with the equivalent substructure in either **a,** SAM1 or **b,** SAM2. Sticks are colored according to ranking (1 – 5 for panel **a** and I – V for panel **b**) in light green, blue, orange, yellow and dark green respectively. Scores from different docking metrics are shown in Supplementary Table 1. Pose 5 from panel k and pose I from panel l are shown as representative poses in Figure 4d.

**Supplementary Table 1 | ICM Flexible Docking results for (S)-SKI-72.** Scoring of ICM docking parameters for 5 binding conformations of (S)-SKI-72 starting aligned with SAM1 adenine and 5 binding conformations of (S)-SKI-72 starting aligned with SAM2 adenine. Each row is shaded the same colour as that used for the described binding pose in Supplementary fig. 15 a (SAM 1 site poses, 1-5) or 15 b (SAM 2 site poses, I-V).

| No. | RTCNN Score | LE Score | ICM-VLS Score | Ligand strain | Van der Waals | Electrostatics | H-bond energy | Hydrophobic energy | Desolvation |
|---|---|---|---|---|---|---|---|---|---|
| **Binding poses starting from SAM1 site (Extended data figure 10k)** | | | | | | | | | |
| **1*** | -31.07 | -10.06 | -25.03 | 14.98 | -46.89 | 25.71 | -12.2 | -10.14 | 43.13 |
| **2** | -30.80 | -13.29 | -21.36 | 8.07 | -50.60 | 30.99 | -11.59 | -10.92 | 46.96 |
| **3** | -30.77 | -11.52 | -23.27 | 11.75 | -52.31 | 29.66 | -11.08 | -10.81 | 46.04 |
| **4** | -25.47 | -3.99 | -16.74 | 12.75 | -56.37 | 37.22 | -9.425 | -11.35 | 48.42 |
| **5** | -26.88 | -13.76 | -28.17 | 14.41 | -45.61 | 24.60 | -14.8 | -10.01 | 46.86 |
| **Binding poses starting from SAM2 site (Extended data figure 10l)** | | | | | | | | | |
| **I** | -29.09 | -7.18 | -18.03 | 10.85 | -50.29 | 33.15 | -10.36 | -10.61 | 44.67 |
| **II** | -25.78 | 4.25 | -6.58 | 10.83 | -51.08 | 32.74 | -6.12 | -10.99 | 47.39 |
| **III** | -27.17 | -8.77 | -14.68 | 5.91 | -44.74 | 23.13 | -8.49 | -9.50 | 43.26 |
| **IV** | -31.38 | -12.96 | -19.45 | 6.49 | -50.65 | 33.12 | -11.45 | -10.93 | 47.24 |
| **V*** | -26.35 | -18.95 | -28.73 | 9.78 | -50.39 | 28.45 | -13.34 | -10.35 | 44.50 |

*Poses shown in Figure 4d

**Supplementary Table 2 | gRNA and ssODNs used for CRISR/Cas9 homology-directed repair.** Parts of the gRNA are highlighted in green, PAM sites in purple, codons of interest are underlined and introduced mutations are displayed in red (desired mutations and PAM site mutations).

| | gRNA (5' – 3') | ssODN (5' – 3') |
|---|---|---|
| R388Q | ACGAGTTCCCTAACGGCCGC | GTATTTGCAAGGAAGGTCTGCAGGCCCTCACCATTGGCCGTTAGGGAACTCGTCCCACTCCTGGGTACGGTAGATGTAACTCTTTGGTCTGGAGGCCCAGAAGATGGGACGTACATCTTCCTCT |
| T573V | CTGACGGGATCCACTACGGT | GGCATCTTCCCTGGGCGAGAGATCATCCAGCCCGTCGTAGTGGATCCCGTCAGCTTCATGTTCTGGAAGGTAAAGGAGCCGGGGGCAAGCTTGCCCCGCCCACCTGGAAAACCGTGGGGAGGGA |

**Supplementary Table 3 | Primers used in site-directed mutagenesis experiments.**

| Mutation | | Primer (5´ --> 3´) |
|---|---|---|
| Trp583Ala | F | CCCGTCAGCTTCATGTTCGCGAAGGACGAGGCC |
| Trp583Ala | R | GGCAAAGGCCTCGTCCTTCGCGAACATGAAGC |
| Trp389Ala | F | GAGTTCCCTAACGGCCGCGCGGGCAATTCCTC |
| Trp389Ala | R | AGGGGAAGAGGAATTGCCCGCGCGGCCGTTAGG |
| Asn386Ala | F | GAGTGGGACGAGTTCCCTGCCGGCCGCTGGG |
| Asn386Ala | R | GGAATTGCCCCAGCGGCCGGCAGGGAACTCGTC |
| Phe384Ala | F | ACCCAGGAGTGGGACGAGGCCCCTAACGGCCGC |
| Phe384Ala | R | GCCCCAGCGGCCGTTAGGGGCCTCGTCCCACTC |
| Tyr404Ala | F | GGGGAGCTGAAGGACTACGCCCTCTTCTACCTG |
| Tyr404Ala | R | GCTCTTCAGGTAGAAGAGGGCGTAGTCCTTCAG |
| Thr573Ala | F | CGAGAGATCATCCAGCCCGCCGTAGTGGATCCC |
| Thr573Ala | R | GCTGACGGGATCCACTACGGCGGGCTGGATGAT |
| Thr573Val | F | CGAGAGATCATCCAGCCCGTCGTAGTGGATCCC |
| Thr573Val | R | GCTGACGGGATCCACTACGACGGGCTGGATGAT |
| Glu463Gln | F | GATGAGCCCCTGGCGGCTCAGACCAGCCTGCTG |
| Glu463Gln | R | CTCCTTCAGCAGGCTGGTCTGAGCCGCCAGGGG |
| Arg388Ala | F | GACGAGTTCCCTAACGGCGCCTGGGGCAATTCC |
| Arg388Ala | R | GGAAGAGGAATTGCCCCAGGCGCCGTTAGGGAA |
| Arg388Gln | F | GACGAGTTCCCTAACGGCCAATGGGGCAATTCC |
| Arg388Gln | R | GGAAGAGGAATTGCCCCATTGGCCGTTAGGGAA |
| Arg388Glu | F | GACGAGTTCCCTAACGGCGAATGGGGCAATTCC |
| Arg388Glu | R | GGAAGAGGAATTGCCCCATTCGCCGTTAGGGAA |
| Gln485Ala | F | ATCCTCACCATCAACTCAGCGCCCAACATCAAC |
| Gln485Ala | R | CTTCCCGTTGATGTTGGGCGCTGAGTTGATGGT |
| Gln485Leu | F | ATCCTCACCATCAACTCACTGCCCAACATCAAC |
| Gln485Leu | R | CTTCCCGTTGATGTTGGGCAGTGAGTTGATGGT |
